# Meta-Research: A Poor Research Landscape Hinders the Progression of Knowledge and Treatment of Reproductive Diseases

**DOI:** 10.1101/2021.11.16.468787

**Authors:** Natalie D Mercuri, Brian J Cox

## Abstract

Reproductive diseases have gone under the radar for many years, resulting in insufficient diagnostics and treatments. Infertility rates are rising, preeclampsia claims over 70 000 maternal and 500 000 neonatal lives globally per year, and endometriosis affects 10% of all reproductive-aged women but is often undiagnosed for many years. Changes in policy have been enacted to mitigate the gender inequality in research investigators and subjects of medical research. However, the disparities in reproductive research advancement still exist. Here, we analyzed the reproductive science research landscape in attempt to quantify the gravity of the current situation. We find that non-reproductive organs are annually researched 5-20 times more than reproductive organs leading to an exponentially increasing relative knowledge gap in reproductive sciences. Additionally, reproductive organs (breast and prostate) are mainly researched when there is a disease-focus, leading to a lack of basic understanding of the reproductive organs. This gap in knowledge affects reproductive syndromes, as well as other bodily systems and research areas, such as cancer biology and regenerative medicine. Action must be taken by current researchers, funding organizations, and educators to combat this longstanding disregard of reproductive science.

## INTRODUCTION

Biological reproduction is essential to the continuation of a species. Despite the continued growth of the global human population, infertility rates are rising. More critically, adverse pregnancies that include preterm delivery, low birthweight, hypertensive and gestationally diabetic impact the acute and chronic health of the population. In fact, 20% of all pregnancies require medical intervention and in lower resource settings, pregnancy and delivery complications account for over 90% of all maternal and neonatal deaths(WHO, 2019). In 1992, the Institute of Medicine (United States (US)) Committee on Research Capabilities of Academic Departments of Obstetrics and Gynecology published and outlined areas needing improvement, such as low birthweight infants, infertility, and pregnancy-induced hypertension (Townsend, 1992). Two decades later, and despite the essential nature and impact of the reproductive system, these issues are still major challenges in reproductive health.

Infertility is a global health problem that affects approximately 8-12% of all couples (Inhorn & Patrizio, 2015). The rates of infertility have increased since 1990 and continues to rise in prevalence worldwide (Mascarenhas et al., 2012; Sun H et al., 2019). Beyond emotional complications of infertility, there are real physiological consequences for many people. Polycystic ovarian syndrome (PCOS) is the most common endocrine-metabolic disorder affecting females of reproductive age, with a prevalence of approximately 8% in the US (Bednarska & Siejka, 2017; Trivax & Azziz, 2007). The various hormonal disturbances in PCOS predispose these individuals to multiple comorbidities, such as cardiovascular disease (CVD), diabetes, and metabolic syndrome. The metabolic disorder has been well-established, with evidence of insulin resistance and hyperinsulinemia in 60-80% of all PCOS patients (Lazaridou et al., 2017). Research in PCOS has focused on managing symptoms, rather than uncovering the underlying pathogenesis to discover more clear diagnostics and targeted treatments. Endometriosis is a disease characterized by the growth of endometrial cells–the lining of the uterus–outside the uterine cavity that affects an estimated 10% of all reproductive-aged women (Chapron et al., 2019; Kiesel & Sourouni, 2019). Endometrial cell growth outside the uterine cavity may result in severe pelvic pain, possible infertility, and intense dysmenorrhea–painful menstruation (Mehedintu et al., 2014). However, cases of endometriosis may present asymptomatically, which contributes to the difficulty in diagnosis and may result in individuals being undiagnosed for 8-12 years (Kiesel & Sourouni, 2019; Mehedintu et al., 2014). Undiagnosed endometriosis may induce damage and dysfunction of surrounding organs that increases risk of other comorbidities, such as cancer (Kvaskoff et al., 2015).

Pregnancy is also complicated by different pathologies. Global low birthweight is highly correlated to neonatal demise. Between 2000 and 2015, the low birthweight rates were reduced minimally (3%) and are not on track to meet the World Health Organization reduction goal of 30% by 2025 (Blencowe et al., 2019). Preeclampsia is a specific pregnancy-related hypertensive disorder that occurs in approximately 4-10% of all pregnancies, varying by country and region (Fox et al., 2019; Rana et al., 2019). While induced pre-term delivery has reduced the acute health risk, there are still over 70 000 maternal and 500 000 fetal deaths globally per year (Rana et al., 2019). Preeclampsia mortality rates are higher in developing countries due to a lack of access to early healthcare required for the treatment of these individuals; whereas, developed countries face higher economic burdens because of the short- and long-term care needed (Ghulmiyyah & Sibai, 2012).

Of those that survive preeclampsia and induced pre-term delivery, mothers have a higher risk of developing CVD and renal disease, and neonates are at a higher risk of obesity and mental health disorders (Bale et al., 2010; Williams, 2011). In the US, the healthcare cost of preeclampsia within the first year of delivery alone is over $2 billion for both the mother and child (Stevens et al., 2017). Pre-term delivery management with and without preeclampsia contributes a healthcare burden of $26 billion per year (Behrman et al., 2007), not including additional long-term cost associated with chronic health problems of survivors.

Despite the prevalence and health impact of reproductive diseases, they have not received similar levels of research attention compared to other bodily systems. Consequently, compared to treatment advances for cancers and heart disease, reproductive pathologies are often difficult to diagnose, properly treat, and at increased risk of comorbidity development. From the literature, there is a long-understood deficit in reproductive basic and applied research. A lack of progress in this area means a continued and increasing acute and chronic health care burden caused by reproductive pathologies. Research deficits in reproduction may result from historic and ongoing systemic cases against female-focused research and political and legal challenges to female reproductive health. Here, we aim to quantify and analyze the delayed advancement of reproductive research in comparison to other biological disciplines. Within this exploratory analysis, we sought to identify possible causes of the continued knowledge gap alongside potential solutions to close this knowledge gap.

## RESULTS

### Total Matching Articles

Our first goal was to benchmark published research on major organs, including the reproductive organs, to quantify research output. Twelve organs of reproductive and non-reproductive systems were chosen for comparison and publication data was extracted from the National Center for Biotechnology Information’s (NCBI) PubMed database (Table I.). This analysis revealed that the total number of articles on non-reproductive organs were 5-20 times higher compared to reproductive organs. This was true for both female- (Breast, Ovary, Uterus, Placenta) and male- (Testes, Prostate, Penis) specific reproductive organs.

**Table I.** Article keyword and total number of matching results from

| Keyword | Total Matching Articles |
| --- | --- |
| Brain | 1 024 560 |
| Heart | 857 851 |
| Liver | 837 791 |
| Lung | 628 410 |
| Kidney | 444 876 |
| Breast | 443 590 |
| Prostate | 190 916 |
| Ovary | 86 000 |
| Uterus | 68 024 |
| Placenta | 58 776 |
| Testes | 33 069 |
| Penis | 15 738 |

### Publication Rates Over Time

The research landscape may change over time, and recent increased efforts to remove gender bias in research may improve reproductive research output. We reassessed the publications on the 12 organs as a function of time from the years 1915–2020 (Figure 1.a). Publication rates of all organs are relatively low with a minimal rate of increase until the 1940s. Following the 1940s, the non-reproductive organs display a near-uniform rise in publications until the early 1970s where the publication rates begin to diverge and increase rapidly. Throughout this rise in organ-focused research, the reproductive organs do not follow the same pattern. The majority of reproductive organs examined maintain a very low publication rate through to 2020. Breast and prostate were the only reproductive organs to increase in publication to a rate similar to the kidney, the least studied non-reproductive organ in our list. To investigate why breast and prostate were the only two reproductive organs showing similar rates to non-reproductive organs, subsequent analysis was conducted on the effects of disease-driven applied research versus basic research.

**Figure 1.**
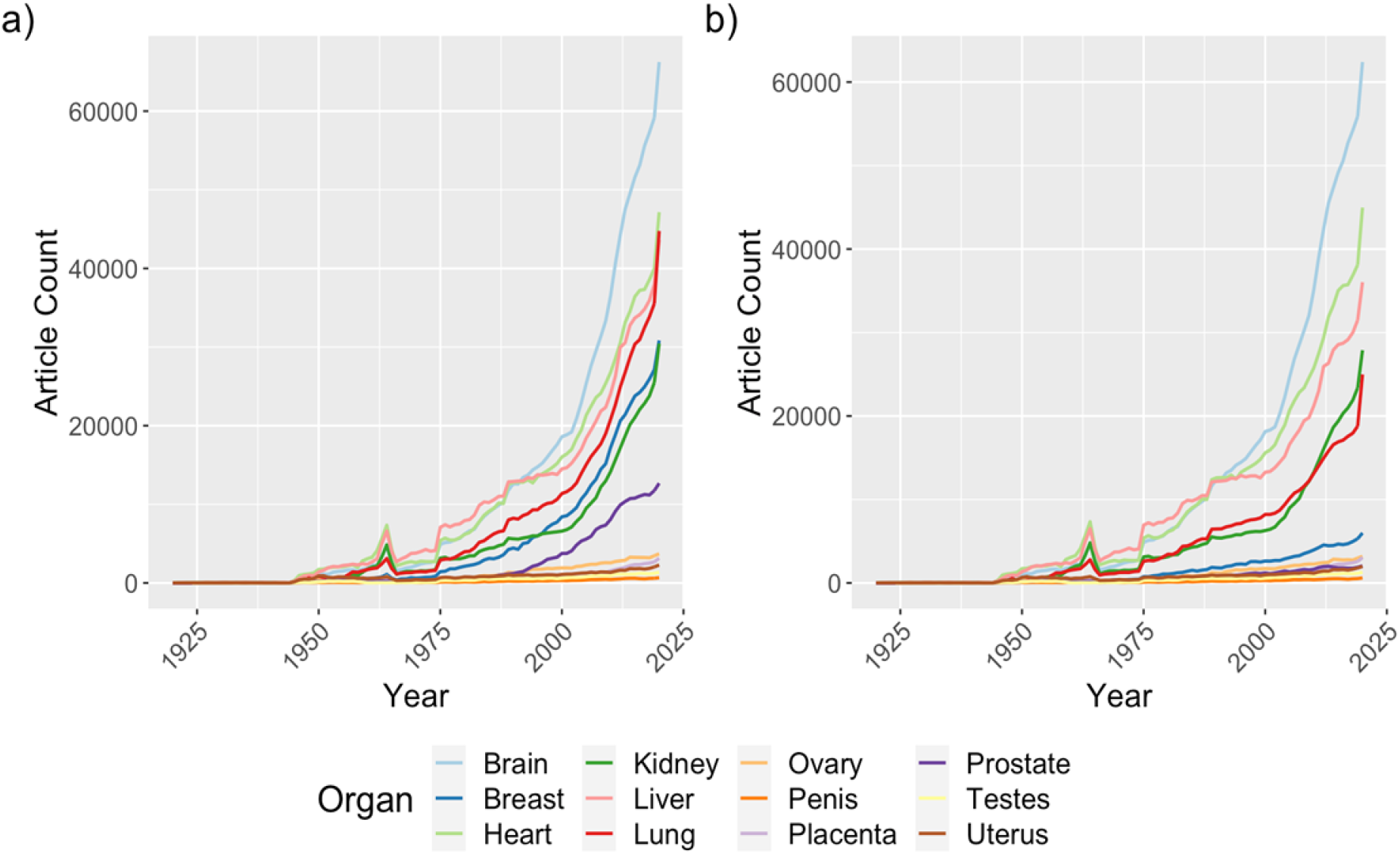
a) Publication rates for different organs as keywords in the title/abstract from PubMed. b) Publication rates for different organs as keywords with search parameter "NOT Cancer" in the title/abstract from PubMed. The years 1915–2020 are represented. Substantial declines in prostate and breast pubication rates are observed.

### Basic vs. Applied Research

Beginning in the 1970s, a war on cancer was initiated by the National Institutes of Health (NIH) and both breast and prostate are organs associated with sex-specific cancers. We reassessed publication data with the added search parameter “NOT cancer” to eliminate cancer-based research (Figure 1.b). We observed that while most non-reproductive organ publications show a decrease of approximately 20%, the breast and prostate publication rates are significantly reduced to the level of the other reproductive organs, losing more than 80% of their publications. Our observation of the impact of cancer on the publication rates motivated a broad look at applied versus basic research of published reproductive-focused research versus other organs.

Translational or applied research is best when based on solid understandings of basic biology. As seen in Figure 1b, breast and prostate publication rates are highly influenced by research associated with cancer—accounting for majority of the publications. We sought to calculate a ratio of basic versus applied research to gauge if reproductive and non-reproductive organs are different. Plotting of the basic versus applied research article count revealed an unexpected high degree of variation of the organs (Figure 2.). The non-reproductive organs brain, heart, and liver plot to the far right of the correlation line with a nearly three-fold higher proportion of basic over applied research. The kidney and lung are relatively close to equivalence of basic and applied research ratio. In contrast, breast and prostate are highly skewed to applied research by a factor of three (Figure 2.). The remaining reproductive organs plot biased to basic research by 1.5 to 2-fold, although the overall publications relative to the basic research-enriched organs of brain, heart, and liver are 10 times less. While suggestive that many reproductive organs achieve a good balance of basic versus applied research, the paucity of research is highly problematic to the field.

**Figure 2.**
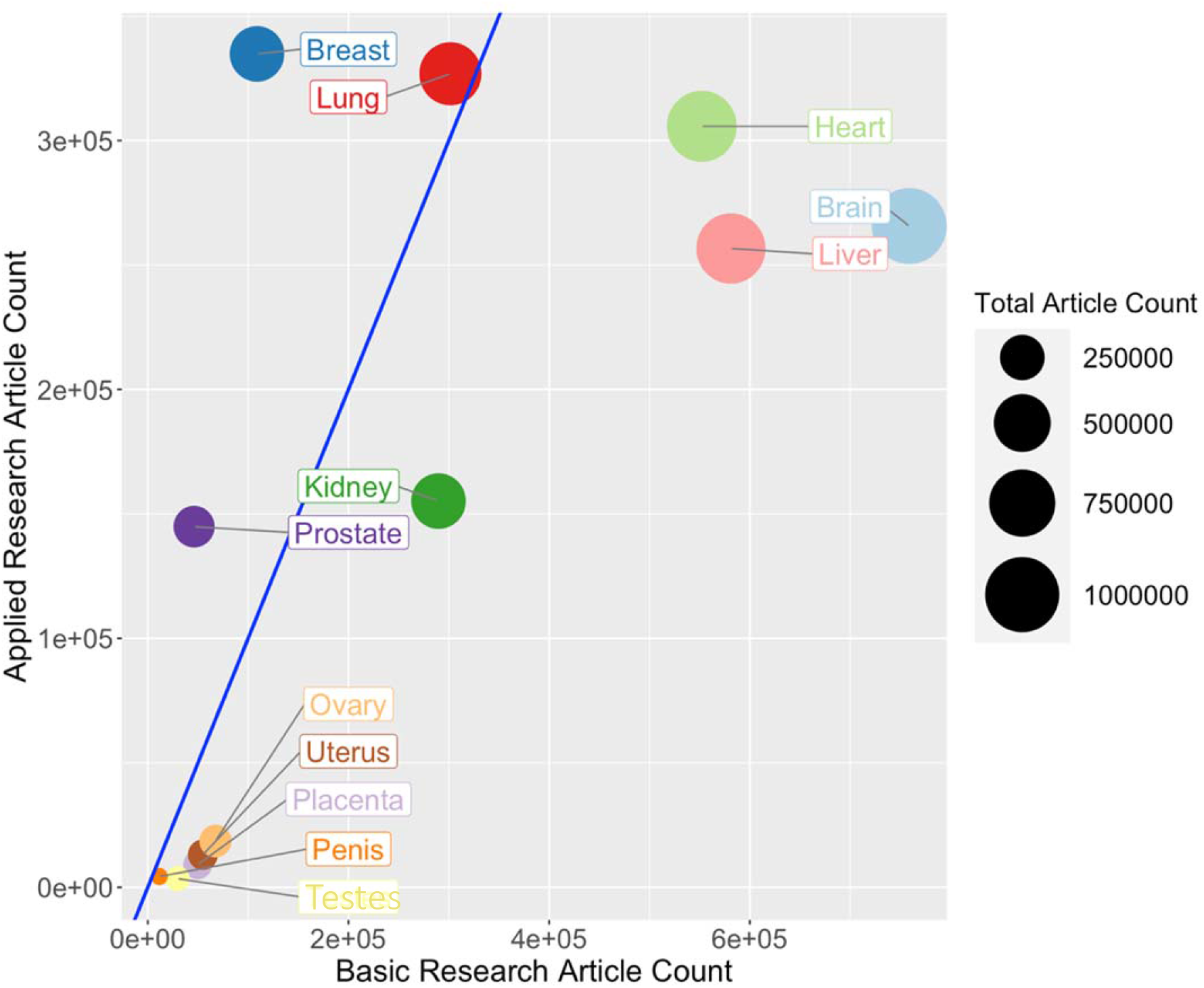
Comparison of PubMed keyword search results of an organ, separated by basic and applied research. Each bubble represents one of the organs and its size represents the total research article count, basic and applied research combined. The blue line is the equal basic and applied counts. Articles in PubMed up to, and including, the year 2020 are represented.

### Comparison of Publication Rates of Diseases by Prevalence

As the majority of research grants are awarded though publicly (tax) funded agencies, an assumption is that disease prevalence and impact (quality of life and mortality) might drive research rates. Acute rates of disease are estimated from the yearly new case diagnoses. Considering acute diseases (cancers and preeclampsia), within the USA, there are approximately 248,530 prostate and 281,550 breast cancers diagnoses per year, which represent 0.08% of the total population each year. Preeclampsia represents about 200,000 diagnoses each year in the USA representing 0.06% of the population (Fingar et al., 2017). Publications of research focused on each of these diseases did not reflect the annual new caseloads. Despite similar annual rates of diagnosis, breast cancer publications outnumbered preeclampsia by nearly 10 times. Until 2005, preeclampsia and prostate cancers publication rates were similar, but by 2019, prostate cancer publications outnumbered preeclampsia by two-fold notwithstanding a similar annual prevalence of the disease. Curiously, ovarian cancer with a 10 times fewer new diagnoses per year (USA: 21,000) to preeclampsia also exceeds preeclampsia publications by at least 2:1 (Figure 3.).

**Figure 3.**
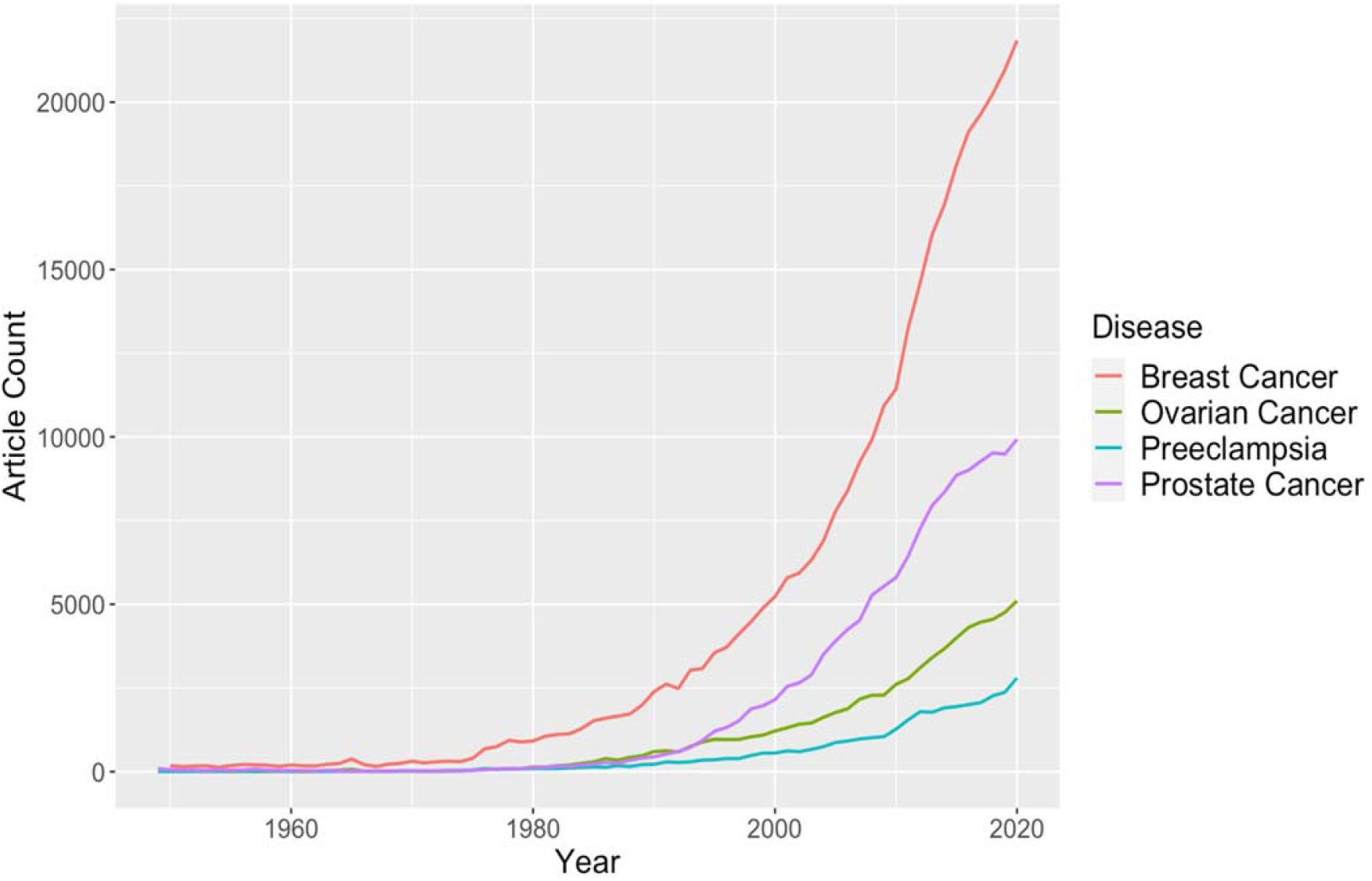
Comparison of publication rates from PubMed of keywords in the title/abstract with an acute pathology focus. The years 1949–2020 are represented. Annual publication counts were extracted for these years from PubMed.

Considering chronic disease burden of the population, Autism Spectrum Disorder (~1%), Crohn’s Disease (~0.24%), Lupus (~0.5%), and tuberculosis (~2%) (CDC; Crohn’s and Colitis Foundation of America; The Lupus Foundation of America; Langer *et al.*, 2019). PCOS is a prevalent reproductive disease affecting nearly 8% of women of reproductive age, which translates into 1.8% of the population (Trivax & Azziz, 2007). Despite a similar prevalence of PCOS to Autism and tuberculosis, PCOS-focused research publications were outnumbered 4-5 times by the year 2020 (Figure 4.). While even diseases of lower prevalence relative to PCOS (Crohn’s and Lupus) also outnumbered PCOS publications at least 2-fold in the year 2020.

**Figure 4.**
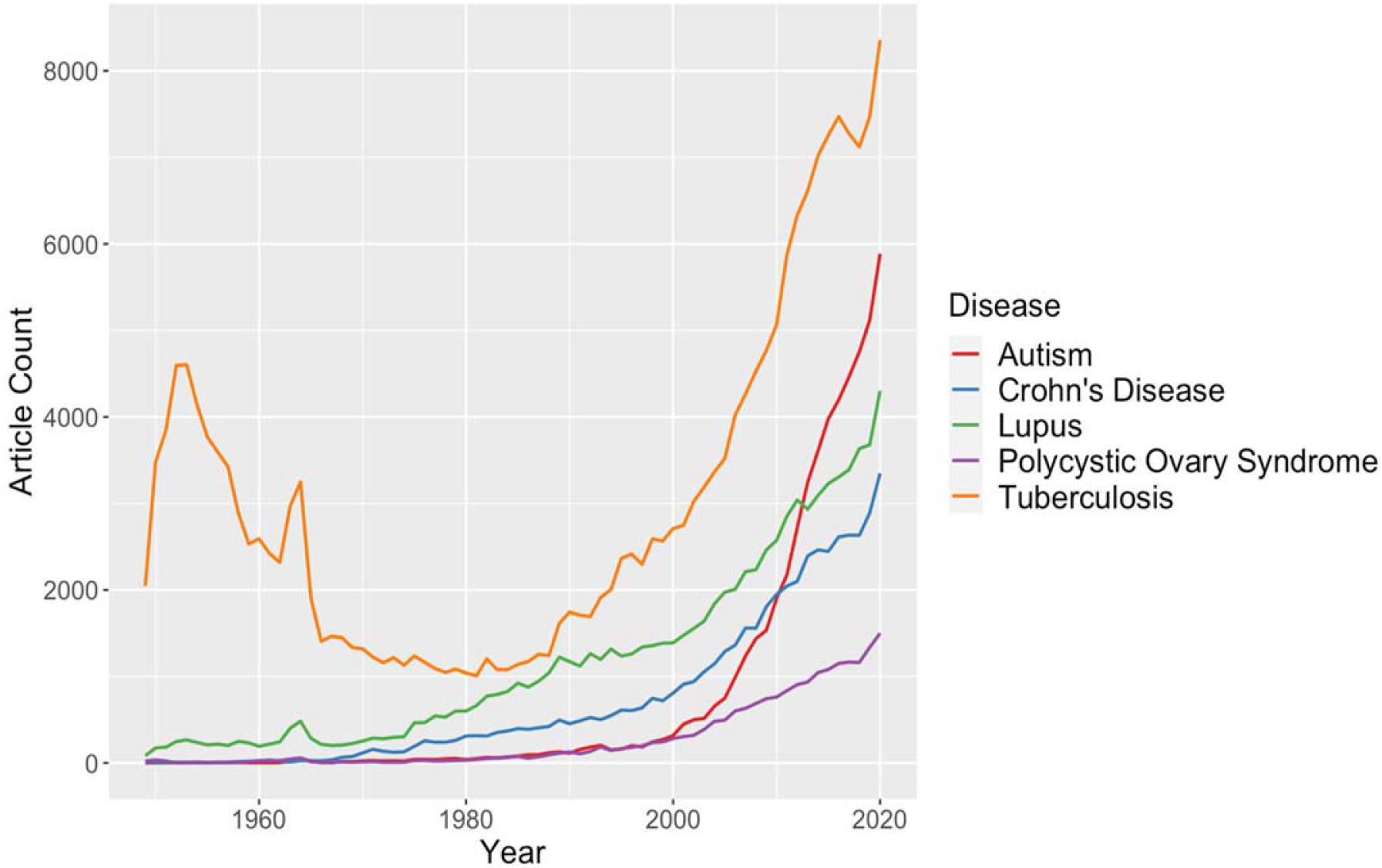
Comparison of publication rates from PubMed of keywords in the title/abstract with a chronic pathology focus. The years 1949–2020 are represented. Annual publication counts were extracted for these years from PubMed.

### Research funding and reproductive research

To investigate the funding trends of different areas, the Canadian Institutes of Health Research (CIHR) and NIH funding databases were used. The keywords brain, heart, liver, breast, prostate, placenta, and testes were entered into each database and funding data from the years 2013–2018 were extracted. Table II (CIHR) and Table III (NIH) display the number of projects for each keyword and the corresponding average funding amount per grant. Our analysis found that the mean grant amounts for the CIHR and NIH are similar between different keyword research topics (CIHR: $ 366 000 per grant ± $ 50 000; NIH: $ 493 000 ± $ 50 000). The similar levels of funding amounts between different disciplines are encouraging and may be a result of standard funding guidelines for biomedical research. However, our analysis found that the annual number of funded projects are much higher for non-reproductive organs compared to reproductive organs in both the CIHR and NIH database analysis. An analysis of funded grants at CIHR and NIH using pathology keywords as used for publication analyses (Figure 3 and 4) produced similar finding of low project numbers for diseases with a reproductive focus that are in line with publication rate differences of 2 to 20-fold (Data not shown).

**Table II.** Average funding amount in CDN ($) from CIHR and the number of projects for the keyword from the years

| **Keyword** | **Number of Projects** | **Average Funding Amount (CDN)** |
| --- | --- | --- |
| Brain | 1875 | \$ 415 000 |
| Liver | 1868 | \$ 319 000 |
| Heart | 1398 | \$ 388 000 |
| Breast | 595 | \$ 361 000 |
| Prostate | 137 | \$ 306 000 |
| Placenta | 115 | \$ 399 000 |
| Testes | 12 | \$ 372 000 |

**Table III.** Average funding amount in USD ($) from NIH and the number of projects for the keyword from the years 2013–

|  |  |  |
| --- | --- | --- |
| Keyword | Number of Projects | Average Funding Amount (USD) |
| Brain | 97 886 | \$ 437 000 |
| Heart | 58 863 | \$ 508 000 |
| Liver | 22 898 | \$ 472 000 |
| Breast | 19 147 | \$ 525 000 |
| Prostate | 8 954 | \$ 514 000 |
| Placenta | 2 567 | \$ 513 000 |
| Testes | 366 | \$ 483 000 |

## DISCUSSION

### Overview of Research Landscape

Reproductive organ research is substantially less published than other organs. Our hypothesis was that hindrance in research progression specific to female reproductive organs is due to sex bias. Gender inequality and bias has been an issue since the onset of the fields of biological and medical research. There has been progress in developing policies to increase the representation of women in research (as both subjects and conductors), as well as in providing education on gender inequality for all researchers. Since 1955, there has been an increase in the number of female authors and their publications. However, there is still a low proportion of female representation within the STEM disciplines (Huang et al., 2020). Additionally, the inclusion of females as subjects in clinical research was not standard practice until after 1986 (Liu & DiPietro Mager, 2016). Therefore, there should be an improving trend over time in female reproductive organ research.

Disappointingly, we do not observe a correlation of reproductive research publications and changes in policies and promotion of women in science. Around the 1940s, there is an increase in all research that coincides with the time of World War II, followed by a large spike around the 1970s (Quirke & Gaudillière, 2008). In 1945, Vannevar Bush wrote a letter to the US President that played a vital role in kickstarting basic scientific research and allowed for the progression into the rapid publication rates observed in my analysis in the last two decades (Figure 1.) (Córdova, 2020). However, research on reproductive organs did not increase at the same rates suggesting at worst, a purposeful exclusion and at best, ignorance. However, our analysis also found low publication rates and numbers of funded projects for both female and male reproductive organs relative to non-reproductive organs. This suggests a broader bias against biological and clinical reproductive research in general. Our observations suggest policies alone are not sufficient to overcome the systemic biases in research disciplines and prevent reproductive research from falling further behind.

Cancer research that drove publication rate increases in breast and prostate did not increase basic research on these organs (Figure 1.b) and suggest an interest in the organs only as it pertained to cancer with no knock-on effect to related research areas. Improved diagnostic and awareness likely led to the increase in publications, such as the Breast Cancer Awareness Month (Jacobsen & Jacobsen, 2011) and the prostate specific antigen test screening (Dickinson et al., 2016). The increase in cancer-focused reproductive organ research likely arises from disease-focused patient and foundation advocacy groups. An important consideration is that a lack of basic biology research in reproductive organs may negatively impact progress in diagnostic and treatment discovery of organ-specific pathologies, including cancers (Figure 5.).

**Figure 5.**
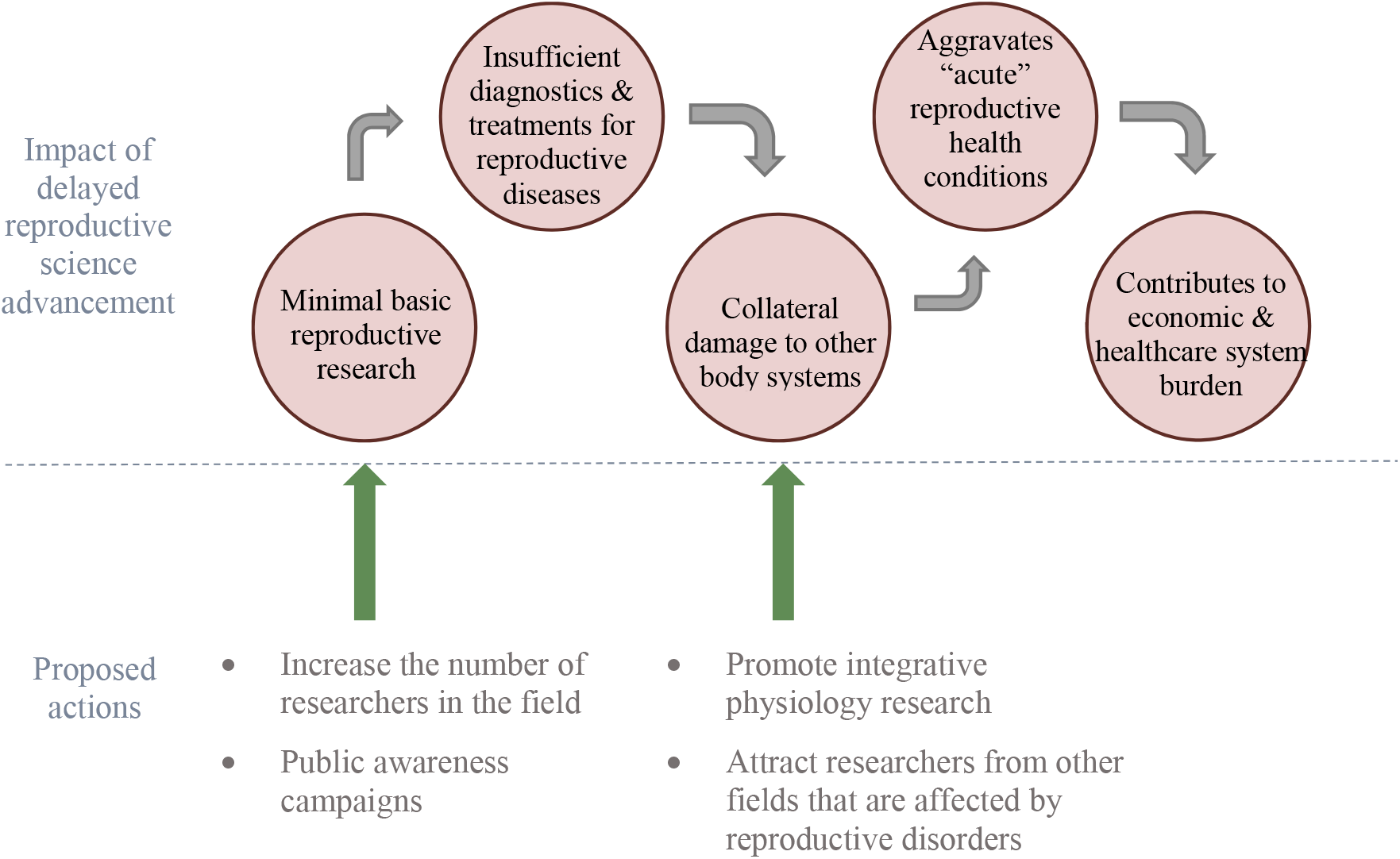
Summary of the problems in reproductive science research reviewed in this article and possible action items as solutions moving forward.

### Basic Research and Funding Trends

Basic science research refers to the exploration of gaps in knowledge, whereas applied research refers to findings that can be translated to create a practical diagnostic or treatment. A lack of basic reproductive science research was cited in the 1992 book “Strengthening Research in Academic OB/GYN Departments” (Townsend, 1992). Our analysis indicates that basic research is a focus for reproductive organs with a strong bias away from applied. Therefore, the likely critical problem to the reproductive field is the overall low level of published research relative to other organs (brain, heart and liver). The low body of knowledge represented by the publication record creates a gap in knowledge that hinders and slows discovery of effective diagnostic and treatment options for pathologies such as preeclampsia, PCOS, and endometriosis. A caveat to our approach is that inconsistencies in data reporting challenge the separation of basic and applied research by keywords alone. However, the discrepancies between these different organs are immense and unlikely to be a result of missing search terms.

In a competitive funding system, publications are correlated to successful grants and dollar values awarded. Across research areas we found that the mean grant dollar amounts per project are similar. However, reproductive organ focused grants were awarded 2-10-fold fewer projects per year – in-line with observed publication rates (Table 1). Critically, as the number and keywords of failed grant applications are not accessible, we could not determine if lower numbers of reproductive grants were due to proportionally lower total application numbers or a bias against funding reproductive research. A possible source of higher failure of reproductive focused grants is the high emphasis placed on applied research, such as for cancer of reproductive organs (breast and prostate), that may arise from a view of basic research as ineffective and inferior compared to applied research (Lee, 2019). Basic research can provide insights into processes that may not be obvious or the intent of the research. Lee encapsulates the gravity of this issue by writing, “basic scientific research ensures we are equipped to deal with issues beyond the limits of our present-day imagination, should they arise.” (Lee, 2019). With a high deficit of basic knowledge on reproductive organs relative to other organ systems, the reproductive medicine field is challenged to describe translational research paths. Granting agencies should examine if funding biases exist. However, generally we suggest that here is a paucity of researchers to enable expanding research on reproductive organs.

### Causes of Reproductive Science Neglect

The possible sources of neglect of the reproductive science are numerous. Overall, there are a variety of stigmas or “taboos” surrounding any topic relating to reproductive function. Menstruation is one function that has faced stigmatization that persists today (Litman, 2018; Pickering, 2019), with women often feeling too embarrassed to talk about this natural process or even complete an essential task, such as purchasing menstrual products at a local store. Political power highly affects reproductive health care and rights over other biological processes. Within many countries, there are ongoing political and legal battles that directly affect access to safe reproductive health care, including contraception, safe abortion, and gender identity rights (Pugh, 2019). Similar struggles around reproductive and sexual health were overcome by decades of progressive action. An example is the Acquired Immunodeficiency Syndrome (AIDS) epidemic in the 1980s. The long delay in recognizing AIDS as a major health issue and implementing research policies perpetuated false ideas surrounding the lifestyles of those affected by the disease and created a barrier to expanding sexual education and seeking healthcare, likely costing many lives (Francis, 2012). Despite great advances in AIDS research and treatment, including social awareness, a public health stigma still lingers in society (Turan et al., 2017). Similar increases in advocacy and public awareness are needed to overcome these barriers for the reproductive sciences.

### Moving forward

Considering our observations from the literature we propose ways to address the research gap (Figure 5.) at a pace that will enable the field to catch up to other areas of biology. Based on our analysis, we need to increase the number of researchers working on reproductive organs and related pathologies. Recent efforts by the NIH, such as the Human Placenta Project (Guttmacher et al., 2014), indicate a recognition of the need to increase research capacity in the area of reproductive sciences and may lead to longer term increases in interest and research capacity.

New researchers may avoid investigations of reproductive biology and disease due to negative perceptions caused by political controversy, low publication levels and poor grant funding numbers. Additionally, the increasing disparity between rates of reproductive biology and disease publication and other biomedical research areas results in reduced exposure of undergraduate and graduate trainees to the reproductive field, further aggravating attempts at increasing the size of the research base. While continued advocacy, education, and political lobbying may overcome many aspects of the social stigma of reproductive sciences, the research gap requires other approaches.

To increase researchers and research output, we may learn lessons from the examples of breast and prostate cancers and autism. In each of these cases, research focused on these diseases rose dramatically from a historically low level. While public campaigns played a large role in this increase, large existing pools of researchers and trainees in the cancer field could switch focus to a different cancer type. A conclusion supported by the observation that there was little or no increase in basic biology research of prostate and breast organs.

We suggest that a similar effect may be a challenge for preeclampsia and other reproductive pathologies as there does not exist a large pool of researchers to refocus on a syndrome. To increase research capacity, we should promote integrative physiology research between reproductive organs and systems of other disciplines with a higher research capacity. Interdisciplinary biomedical research is unarguably crucial to achieving comprehensive healthcare. The paucity of reproductive research identified by our analysis strongly implies a poor understanding and integration of the reproductive systems within human physiology. Yet examples exist to show the benefit of an integrated approach that include reproductive sciences. Female sex hormones are protective against many diseases of aging, such as cardiovascular and neurological, leading to prescription of hormone replacement therapies after menopause in some females (Paciuc, 2020). To improve reproductive sciences research, we should focus on promoting awareness of knowledge gaps to researchers of disciplines outside of the reproductive fields (Figure 5.).

During pregnancy, there are dramatic changes in maternal physiology, including metabolism, the immune system, and cardio-pulmonary systems and consequently, these are the same systems affected by reproductive pathologies. Preeclampsia predisposes the mother to a long-term cardiovascular risk of developing peripheral artery disease, coronary artery disease, and congestive heart failure (Rana et al., 2019). The rising incidence of preeclampsia is directly contributing to the global burden of CVD. Additionally, complications of the liver and kidney are associated with preeclampsia. PCOS and endometriosis are associated with problems in metabolism as well and risk of cancer development. Children born from pregnancies affected by preeclampsia or fetal growth restriction are at a 2.5 times higher risk of developing hypertension and requiring anti-hypertensive medications as an adult (Ferreira et al., 2009; Fox et al., 2019).

The pathological interaction of reproductive with non-reproductive systems and organs should attract investigators from cardiovascular, nephrology and hepatology fields, where there exist 10-20 times the total number of researchers than in reproductive biology. If 1% of the researchers in the cardiovascular field were to refocus on pregnancy related cardiovascular adaptations and pathologies, this would increase reproductive research by 10%.

Our ignorance of the placenta and reproductive biology is impeding other areas of biomedical research. In cancer research, the methylation patterns of tumours look most like those found in the placenta, but why placenta methylation patterns are so unlike all other organs is not known (Rousseaux et al., 2013). In regenerative medicine research, the immune-modulating genes used by the placenta (Szekeres-Bartho, 2002) are being repurposed to generate universally transplantable stem cells and tissues (Han et al., 2019). A poor understanding of reproductive biology is dangerous in consideration of emerging diseases that affect pregnancy and fetal development, such as the recent Zika virus outbreak (Calvet et al., 2016; Schuler-Faccini et al., 2016). There are likely many other broad benefits to better understanding reproductive biology. The time to act is now as waiting longer will not improve the situation.

## METHODS

### Publication Rates

Published research manuscripts were searched in NCBI’s PubMed database (https://pubmed.ncbi.nlm.nih.gov/) as of January 2021, therefore, the entire year of 2020 was accounted for. Keywords for each search pertained to a specific organ or disease and were limited to the title/abstract of the manuscripts. The organs used for these analyses were brain, heart, brain liver, lung, kidney, breast, prostate, ovary, uterus, penis, and placenta. The organ publication timelines were restricted to the years 1915–2020 and the annual article count was extracted. The organ publication timeline was reconducted with the addition of the search parameter “NOT cancer”. The acute and chronic diseases used for this analysis were breast cancer, ovarian cancer, prostate cancer, preeclampsia, PCOS, autism, tuberculosis, Crohn’s disease, and lupus. The keywords for these diseases were: “preeclampsia”, “polycystic, ovary OR ovarian, AND syndrome”, “autism”, “tuberculosis”, “crohn OR crohn’s, AND disease”, and “lupus”. The number of cancer diagnoses were retrieved from the National Cancer Institute (https://seer.cancer.gov/statfacts/). The demographic values for population ages and pregnancy rates were estimated from the US Census Bureau (https://www.census.gov/data/tables/2020/demo/popest/2020-demographic-analysis-tables.html)). The disease publication rate timeline was restricted to the years 1949–2020 and the annual article count was extracted.

### Basic vs. Applied Research

Published manuscripts were searched in NCBI’s PubMed database (https://pubmed.ncbi.nlm.nih.gov/) as of January 2021. Two searches were completed for each organ. The first search was the organ in the title/abstract. The second search was the same organ in the title/abstract with the added search parameters “NOT disease, NOT disorder, NOT condition, NOT cancer”. The second search retrieved articles that were considered as basic research for this analysis. The second search article count was then subtracted from the first search article count to represent the articles deemed as applied research for this analysis.

### Funding Rates

Grant funding data was obtained from the Canadian Institutes of Health Research funding database (https://webapps.cihr-irsc.gc.ca/funding/Search?p_language=E&p_version=CIHR) and the National Institute of Health’s funding reporter database tool (https://reporter.nih.gov). Keywords used for these searches were brain, heart, liver, breast, prostate, placenta, and testes. The years were restricted to 2013–2018. The total number of projects pertaining to each search during this time period was extracted, as well as the total amount of funding for those projects that was then averaged.

### Graphing

All graphs were produced using R (version 4.0.2) in R Studio (version 1.3.1073). R packages used were ggplot2, tidyverse, formattable, gridExtra, RColorBrewer, ggrepel.

## DATA AVAILABILITY

Data underlying this article was extracted from publicly accessible databases. Code used to generate analyses will be made available after reasonable request to the corresponding author.

## ACKNOWLEDGEMENTS

Thank you to the University of Toronto and the Department of Physiology for providing the opportunity and supporting completion of this review.

## AUTHORS’ ROLES

NDM developed the idea, collected the data, designed the tables and figures, and wrote the manuscript. BJC developed the idea, supervised the project, provided expert opinion on the field and feedback on the manuscript.

